# SARS-CoV-2 utilization of ACE2 from different bat species allows for virus entry and replication *in vitro*

**DOI:** 10.1101/2023.04.19.537521

**Authors:** Kelsey Briggs, Ryan Sweeney, David S. Blehert, Erica Spackman, David L. Suarez, Darrell R. Kapczynski

## Abstract

Severe acute respiratory syndrome coronavirus 2 (SARS-Cov-2) is believed to have a zoonotic origin. Bats are a suspected natural host of SARS-CoV-2 because of sequence homology with other bat coronaviruses. Understanding the origin of the virus and determining species susceptibility is essential for managing the transmission potential during a pandemic. In a previous study, we established an *in vitro* animal model of SARS-CoV-2 susceptibility and replication in a non-permissive avian fibroblast cell line (DF1) based on expression of angiotensin-converting enzyme 2 (ACE2) and transmembrane serine protease 2 (TMPRSS2) from different animal species. In this work, we express the ACE2 of seven bat species in DF1 cells and determine their ability to support attachment and replication of the original SARS-CoV-2 Wuhan lineage virus, as well as two variants, Delta and Lambda. We demonstrate that the ACE2 receptor of all seven species: little brown bat (*Myotis lucifugus*), great roundleaf bat (*Hipposideros armiger*), Pearson’s horseshoe bat (*Rhinolophus pearsonii*), greater horseshoe bat (*Rhinolophus ferrumequinum*), Brazilian free-tailed bat (*Tadarida brasiliensis*), Egyptian rousette (*Rousettus aegyptiacus*), and Chinese rufous horseshoe bat (*Rhinolophus sinicus*), made the DF1 cells permissible to the three isolates of SARS-CoV-2. However, the level of virus replication differed between bat species and variant tested. In addition, the Wuhan lineage SARS-CoV-2 virus replicated to higher titers (10^4.5^-10^5.5^ TCID_50_) than either variant virus (10^3.5^-10^4.5^ TCID_50_) on pass 1. Interestingly, all viruses tested grew to higher titers (approximately 10^6^ TCID_50_) when cells expressed the human ACE2 gene compared to bat ACE2. This study provides a practical *in vitro* method for further testing of animal species for potential susceptibility to current and emerging SARS-CoV-2 viruses.

## INTRODUCTION

The on-going COVID-19 pandemic is caused by severe acute respiratory syndrome coronavirus 2 (SARS-CoV-2; SC2). It was first identified in Wuhan, China in 2019 and declared a global pandemic by the World Health Organization (WHO) in March 2020 (1, 2). Symptoms of SC2 typically include fever, chills, shortness of breath, and loss of smell/taste, but severe disease can lead to death. SC2 belongs to the virus family *Coronaviridae*, which are single-stranded, enveloped, positive-sense RNA viruses (3). The family contains several other viruses that cause respiratory illness in humans and animals such as porcine enteric diarrhea (PEDV), infectious bronchitis virus (IBV), and human CoV-NL63 (4–6).

Since first detection of SC2 in humans in 2019, it has rapidly spread across the globe and acquired several mutations giving rise to variants. The variants are categorized into three categories by the WHO: variants of interest (VOI), variants of concern (VOC), and variants under monitoring (VUM) (1). The original Wuhan strain was quickly replaced by a variant containing a D614G mutation in the spike (S) protein. This mutation was associated with increased viral yields in human cells and virus stability (7, 8). Alpha, Beta, and Gamma VOCs followed shortly after with varying degrees of antibody cross neutralization and immunity in humans (1, 9–11). The Delta VOC, which was first detected in India in October 2020, rapidly took over as the dominant variant. This was attributed to immune escape, either from natural infection or vaccination, and increased fitness of the variant to replicate in humans. However, in June 2022, Delta was classified as a “previously circulated” VOC (1, 12–14). Some VOIs include Epsilon, Theta, Mu, and Kappa, which all circulated during 2020-2021, whereas the Lambda VOI circulated until March 2022 (1). The Lambda VOI was shown to be more infectious than previous variants, evade neutralizing antibodies, and have the potential to cause antibody-mediated enhanced disease, all of which contributed to its substantial spread across South America (15). To date, the dominant VOC circulating is the omicron variant and its subsequent lineage. Compared to previous variants, this lineage has a substantial number of mutations throughout the genome, including 30 in the S protein with half of those being in the receptor binding domain (RBD) (1, 16, 17). The origin of Omicron is currently unknown, but it has been postulated that it circulated and adapted in animal reservoirs, then transmitted back to humans (18).

In the past two decades, two other Coronaviruses with high fatality rates in humans, Severe Acute Respiratory Syndrome (SARS) and Middle East Respiratory Syndrome (MERS), were detected for the first time. Both viruses are believed to have originated in bats before disseminating to intermediate hosts and then humans (19–22). Although, to date, the natural host for SC2 is unknown, it is hypothesized to have come from bats (23, 24). Recent studies have shown that bat-borne SC2-like viruses circulate in *Rhinolophus* species in Southeast Asia, but a direct progenitor virus has yet to be found (25). To date, only four bat species have been experimentally infected with SC2, whereas roughly 1400 species of bats are estimated worldwide (26–29). To test susceptibility of every bat species to SC2 would be impractical, but less intrusive and lower-cost methods are available to examine if the virus can replicate within a bat species.

SC2 utilizes angiotensin-converting enzyme 2 (ACE2) as its receptor for host-cell attachment by the S protein (30). Many homologues of ACE2 exist in the animal kingdom; however, mammals have the highest degree of ACE2 conservation, making the potential host range for SC2 extensive (31). Several studies have examined the possible host range based on ACE2 sequence relatedness, but most species have yet to be tested in an *in vitro* or *in vivo* model for SC2 susceptibility (31–33). In this work, we expanded on previous studies looking at species susceptibility to SC2 using a chicken cell line that is nonpermissive to SC2 replication (34). Using transposon mutagenesis, different bat species ACE2 genes were individually inserted into the chicken cell genome and their ability to replicate SC2 and two variants, Delta and Lambda, were assessed. These studies were designed to test whether seven bat species could potentially be a natural or intermediate host for SC2. In addition, they provide an *in vitro* alternative method for testing susceptibility to SC2 in relevant animal species.

## MATERIALS AND METHODS

### Viruses

USA/WA1/2020/Wuhan lineage (BEI NR-52286; Washington strain), USA/PHC658/2021/B.1.617.2 (BEI NR-55611; Delta strain), and Peru/un-CDC-2-4069945/2021/Lineage C.37 (BEI NR-55654; Lambda strain) of SARS-CoV-2 were obtained from BEI Research Resources Repository, National Institute of Allergy and Infectious Diseases, National Institutes of Health (35). Experiments with SC2 were performed in a biosafety level-3 enhanced facility with procedures approved by the U.S. National Poultry Research Center Institutional Biosafety Committee.

### Cell lines

DF1 (avian fibroblast) and Vero (African Green monkey kidney, CCL-81) cells were seeded and propagated with standard procedures for adherent cells, in tissue culture flasks, containing Dulbecco’s Modified Eagle Medium (DMEM) (ThermoFisher Scientific, Waltham, Massachusetts) supplemented with 10% Fetal Bovine Serum (Atlanta Biologics, Atlanta, Georgia) and 1% Antimicrobial-Antimycotic (Gemini-Bio, Sacramento, California). Vero cells were obtained from the International Reagent Resource (FR-243). DF1 cells were cultured at 39℃ whereas Vero cells were cultured at 37℃.

### Construction of plasmids expressing different bat ACE2 genes using the PiggyBac transposon vector

GenBank accession numbers used to construct all bat species plasmids can be found in Table 1. The ACE2 genes from little brown bat (*Myotis lucifugus*), great roundleaf bat (*Hipposideros armiger*), Pearson’s horseshoe bat (*Rhinolophus pearsonii*), greater horseshoe bat (*Rhinolophus ferrumequinum*), Brazilian free-tailed bat (*Tadarida brasiliensis*), Egyptian rousette (*Rousettus aegyptiacus*), and Chinese rufous horseshoe bat (*Rhinolophus sinicus*) were *de novo* synthesized into the PiggyBac® transposon expression plasmids under control of the CMV promoter, expressing EGFP (VectorBuilder Inc., Chicago, Illinois). Frozen *Escherichia coli* plasmid glycerol stocks containing ACE2 were streaked onto Luria-Bertani (LB) agar plates (Invitrogen) containing 100 µg/mL of Carbenicillin (Sigma, St. Louis, Missouri). Single colonies were selected and incubated in 200 mL of LB Broth, containing 100 µg/mL of Carbenicillin, with gentle agitation overnight in an incubator/shaker at 37°C (Amerex Instruments, Concord, California). Plasmid DNA was isolated using ZymoPURE II midiprep plasmid kit (Zymo Research, Irvine, California) per the manufacturer’s protocol.

**Table 1.** Table of ACE2 and TMPRSS2 gene accession numbers. Genbank accession numbers and genes used in this study.

| **Species** | **ACE2** | **TMPRSS2** |
| --- | --- | --- |
| Human | NM_021804.1 | NM_005656.4 |
| Little brown bat (*Myotis lucifugus*) | XM_023753670.1 |  |
| Great roundleaf bat (*Hipposideros armiger*) | XM_019667391.1 |  |
| Pearson's horseshoe bat (*Rhinolophus pearsonii*) | EF569964.1 |  |
| Greater horseshoe bat (*Rhinolophus ferrumequinum*) | AB297479.1 |  |
| Brazilian free-tailed bat (*Tadarida brasiliensis*) | MT663956.1 |  |
| Egyptian rousette (*Rousettus aegyptiacus*) | XM_016118926.2 |  |
| Chinese rufous horseshoe bat (*Rhinolophus sinicus*) | MT394181.1 |  |

### Creation of Transgenic DF1 cells expressing bat ACE2 and human TMPRSS using PiggyBac Plasmid Transposon System

DF1 cells were grown in a T25 flask and transfected with PiggyBac transposon, expressing human TMPRSS2 and an mCherry marker, along with the hyperactive PiggyBac Transposase (hyPBase) utilizing Xfect transfection reagent (Takara-Bio, San Jose, California). Transposase and Transposon DNA were added at 1:5 ratio in Xfect transfection reagent per manufacturer instructions to form the nanoparticle transfection complex. Complete media from cells was replaced with DMEM, and Xfect transfection complex was added for 4-6 hours. After incubation, media containing transfection complex was removed and fresh media containing 10% FBS, 1% P/S DMEM was added. Cells were incubated for 48-72 hours at 39°C 5% CO_2_, after which expression was confirmed using fluorescent microscopy (EVOS M5000). Cells were purified by fluorescence-activated cell sorting (FACS) by gating for mCherry (>99%). FACS-purified DF1 cells containing hTMPRSS2 were grown in a T25 flask and transfected with the PiggyBac transposon system in the same manner as above. Bat ACE2 transposons contained an EGFP marker. Dual transfected cells were then sorted for both GFP and mCherry (>99%). Cells were periodically sorted to enrich the population of GFP- and mCherry-positive cells.

### Fluorescent-activation cell sorting (FACS)

Transgenic cells expressing ACE2 (EGFP), TMPRSS2 (mCherry), or both were grown to 90% confluence, trypsinized, pelleted by centrifugation (1500 RPM for 10 minutes at room temperature), and strained through a 50-µm cell strainer (Thermo Fisher Scientific, Carlsbad, California). Cells were sorted for GFP, mCherry, or both at the University of Georgia (Athens, Georgia) Flow Cytometry Core Center using a Bio-Rad S3e cell sorter (Bio-Rad, Irvine, California).

### RNA extraction and RT-PCR for bat ACE2 and human TMPRSS2 DF1 cell lines

RNA lysates from all cell lines were obtained, including a DF1 (-/-) negative control, as previously described (36). RNA extractions were carried out using the ZYMO Direct-zol Mini-Prep Plus Kit (Zymo Research, Irvine, California) per manufacturer’s instructions.

To check expression levels of ACE2, RT-qPCR was performed. RNA was extracted as previously described, and 7.5 µL of RNA was used per reaction. Human ACE2 primers, universal bat ACE2 primers, and chicken 28S primers were used as previously described (34). Luna® Universal One-Step RT-qPCR Kit (NEB, Ipswich, Massachusetts) was used to quantify the RNA samples per the manufacturer’s guidelines. Values were normalized to the chicken 28S (house-keeping gene) and DF1 (-/-) was used as the negative control to calculate the ΔΔCT values.

### Detection of ACE2 and TMPRSS2 protein expression by immunoblot

Western blot detection was performed as previously described (37). Cells were lysed from a 6-well plate in 2X Laemmli SDS sample buffer plus 2-mercaptoethanol, then boiled at 95℃ for 5 minutes. Samples were resolved on a 10% SDS-PAGE gel, then transferred to a polyvinylidene difluoride (PVDF) membrane (Bio Rad, Irvine, California). The blot was then incubated overnight at 4°C in primary antibody in 2% milk. Primary monoclonal antibodies included mouse anti-human ACE2 (1:1500) (Origene, Rockland, Maryland), rabbit anti-human (1:1000) TMPRSS2 (Abcam, Cambridge, United Kingdom), and mouse anti-beta actin (1:2000) (Invitrogen, Carlsbad, California). The blot was washed 3x in PBST, then incubated for 1 hour at room temperature, in secondary antibody diluted 1:2500 with gentle rocking. Secondary antibodies included Cy3-conjugated goat anti-mouse IgG secondary antibody (Jackson Immuno-Research, West Grove, Pennsylvania), and goat anti-rabbit Dylight™ 594 secondary antibody (ThermoFisher, Carlsbad, California). The blot was washed again in PBST and imaged on a G:Box mini6 (Syngene International Ltd, Bengaluru, India).

### Detection of ACE2 and TMPRSS2 protein expression by immunofluorescence assay

Cells were seeded into an I-Bidi 8-well chambered slide (ThermoFisher, Carlsbad, California) at a density of 4 x 10^4^ in 500 µL DMEM containing 10% FBS, 1% P/S, and grown overnight as above. When cells reached 75% confluence, media was removed and virus was added at MOI of 1. After 48 hours, the media was removed and cells were fixed for 5 minutes at 4°C in 1:1 ice cold ethanol:methanol. Cells were then washed twice with cold PBS. Cells were blocked for one 1 hour at room temperature, then washed 3 times with PBS. The primary antibody, rabbit anti-Spike MAb (Origene, Rockland, Maryland), diluted 1:250, was added for 1 hour at room temperature. Cells were washed 3 times with PBS and incubated in the secondary antibody, goat anti-rabbit IgG H&L (Cy3-conjugated) (Abcam Cambridge, United Kingdom) diluted 1:500 in PBS, for 1 hour at room temperature. Cells were then washed 3 times with PBS and counterstained with DAPI (Invitrogen, Carlsbad, California) for 5 minutes. Immunofluorescence was visualized with an EVOS 5000 (Invitrogen, Carlsbad, California).

### Comparison of SARS-CoV-2 replication dynamics among cell lines

Each cell line (bat ACE2, positive control DF1 (+/+), and negative control DF1 (-/-) was infected with SC2 at an MOI of 1 in a 6-well plate in triplicate. For each cell line, media was removed from three wells and 0.4 mL of virus inoculum was added. Virus preparation was performed as previously reported (34). The same volume of sterile medium was used as a sham inoculated control. The plates were incubated for 1 hour at 37°C, 5% CO_2_. Each well was washed 3 times with sterile PBS prewarmed at 37°C to remove unbound virus. Finally, 3 mL growth medium was added to each well and the plates were incubated at 37°C with 5% CO_2_. Supernatant (0.2 mL) was collected from each well individually at 2, 6, 24, 48 and 72 hours post inoculation (HPI) for detection of replicating virus by RT-PCR. Cytopathic effect was determined by microscopy (EVOS 5000). After 72 HPI, plates were frozen and thawed at −80°C (3x total), and 0.4 mL of cell culture supernatant was transferred onto fresh 6-well plates containing cells for pass 2. The timepoints were repeated to confirm infectious virions were produced in the avian cells.

### Quantitative real-time RT-PCR to detect SARS-CoV-2 replication

Quantitative RT-PCR was utilized to detect and determine virus titers in cell culture supernatants. RNA was extracted with the Ambion Magmax kit (ThermoFisher, Carlsbad, California). The U.S. Centers for Disease Control N1 primers and probe for SARS-CoV-2 were used with the AgPath ID one-step RT-PCR kit (38). The cycling conditions for the RT step were modified to accommodate the recommended kit conditions. A standard curve of RNA from each titrated SC2 virus stock was run in duplicate to establish titer equivalents of virus, and the viral titer was extrapolated from the standard curve.

### ACE2 and TMPRSS2 genetic analysis

ACE2 and TMPRSS2 protein sequences from human and bat species were obtained from GenBank. Sequences were aligned with Geneious Prime (Auckland, New Zealand). A global alignment with free end gaps was performed on available ACE2 and TMPRSS2 sequences. BIosum45 with a threshold of 0 was used for percent similarity.

### Statistical analysis

Viral titers at 48 HPI were compared with the two-way ANOVA with Tukey multiple comparison (Prism 9.1.0 GraphPad Software, San Diego, California). Different lower-case letters indicate statistical significance between compared groups. All statistical tests used p < 0.05 as being statistically significant.

## RESULTS

### Analysis of bat ACE2 genes

We obtained ACE2 protein sequences from seven different bat species for phylogenetic analysis and compared them to other animals previously tested (34). The bat species grouped together, demonstrating that they stemmed from a single ancestral species (Figure 1). Percent similarities ranged from 86-99% between all animals, and 91-99% between bat species (Supplemental Figure 1). Chickens, which are not susceptible to SC2 infection, had the lowest similarity to all other animals and was used as an outgroup for the analysis (28, 39). Interestingly, bat ACE2 proteins were generally as similar to horse, pig, and cat ACE2 as human (93-96%) (Figure 1). However, to date, horses do not appear susceptible to SC2 infection, and *in vitro* studies have corroborated this (34, 40). Several unique areas containing less than 80% similarity were observed in the ACE2 alignment. In particular, amino acid regions 15-22, 91-94, 209-215, 254-258, 608-624, 671-685, and 788-809 had high degrees of variability (Supplemental figure 2).

**Figure 1.**
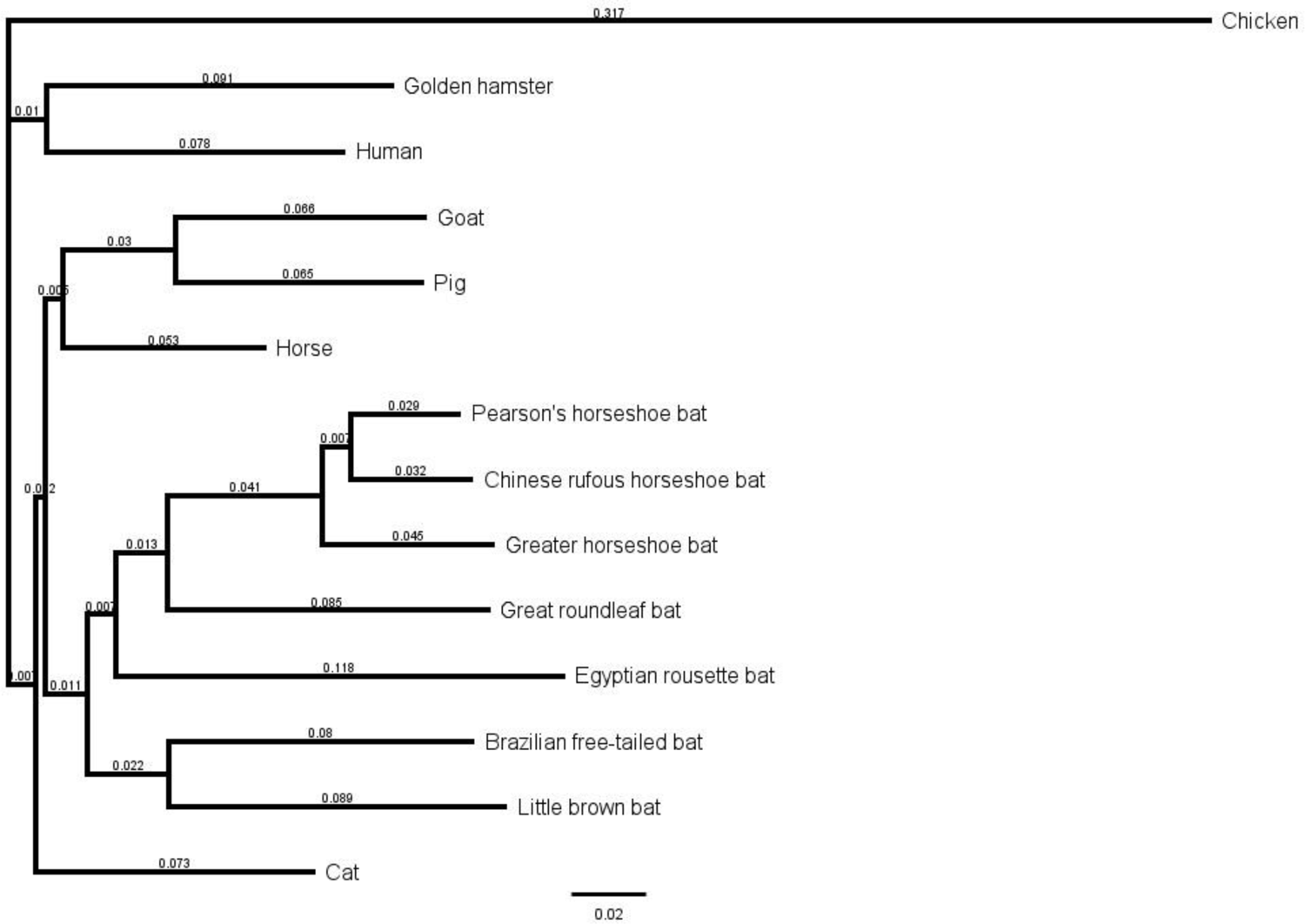
Phylogenetic analysis of ACE2 proteins. Amino acid sequences were obtained from Genbank and aligned using Geneious Prime. A global alignment with free end gaps was performed on the ACE2 sequences. Phylogenetic tree illustrates the relative distances between different ACE2 proteins. Chicken ACE2 was chosen as outgroup because it is not recognized by SC2 spike protein. Distances are labeled and scale bar is shown.

**Figure 2.**
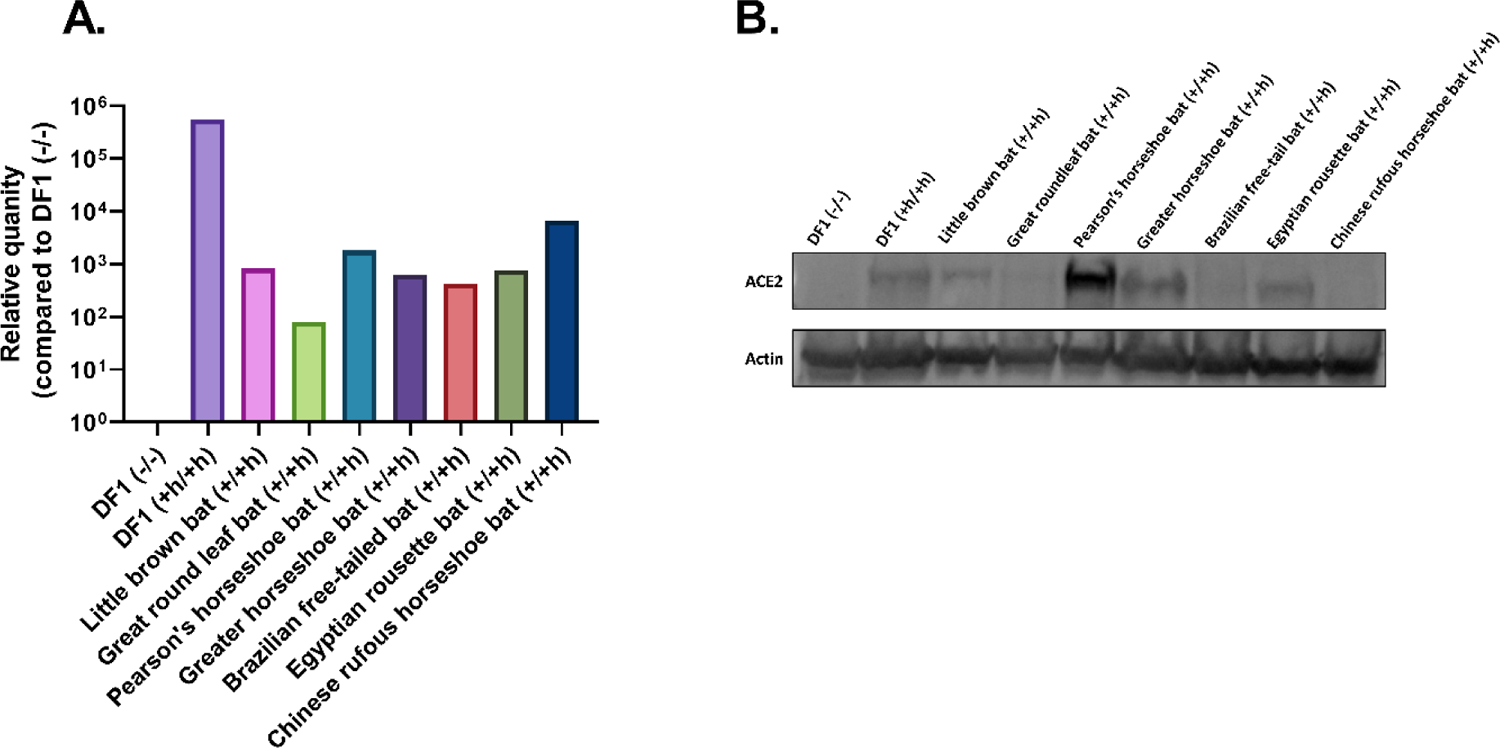
Expression of ACE2 in DF1 cells. DF1 cells lines containing bat ACE2 and human TMPRSS2 were examined for expression of ACE2. **(A)** ACE2 mRNA levels were measured in the cell lines using RT-qPCR. Values were normalized to the chicken 28S housekeeping gene. DF1 (-/-) cells were used as a negative control to calculate the ΔΔCT values. **(B)** ACE2 protein expression was measured by western blot. Cell lysates were resolved by SDS-PAGE and then transferred to a PVDF membrane. The blot was probed with a mouse anti-human ACE2 and a rabbit anti-human actin antibody, followed by a Cy3-conjugated goat anti-mouse IgG and goat anti-rabbit Dylight™ 594 secondary antibody for detection.

### Development of cell lines expressing ACE2 and hTMPRSS2 from different bat species

The human TMPRSS2 (hTMPRSS2) gene was inserted into an avian DF1 cell line using lentivirus delivery, followed by a bat ACE2 gene using the PiggyBac Transposon system. Expression levels of ACE2 and hTMPRSS2 were measured by RT-qPCR (Figure 2 and Supplemental Figure 3). All bat cell lines had at least 100-fold greater ACE2 activity than the unmodified, wildtype DF1 cells (DF1 (-/-)) confirming expression of ACE2 by RT-qPCR (Figure 2A). Additionally, ACE2 protein expression was measured by immunoblot using a human monoclonal ACE2 antibody. Detection of ACE2 protein was variable by immunoblot, and ACE2 from the great roundleaf bat, Brazilian free-tailed bat, and Chinese rufous horseshoe bat did not react with the human ACE2 specific Mab as well as the other bats (Figure 2B), which was likely due to sequence variability at the antibody binding site in ACE2. The wild-type DF1 cells (-/-) did not react with the anti-human ACE2 antibody. These results demonstrate that human and bat ACE2 was expressed in the avian DF1 cells.

### SARS-CoV-2 infection and growth kinetics in cells expressing bat ACE2 and hTMPRSS2

DF1 cells expressing a bat ACE2 and hTMPRSS2 were infected with either Wuhan lineage, Delta, or Lambda SC2, and their viral titers were measured to evaluate growth kinetics. Replication of the Wuhan lineage SC2 was variable in the cell lines. However, it grew to significantly lower titers in the cell lines containing a bat ACE2 compared to the DF1 (+h/+h) control cells, which grew SC2 to >10^6^ TCID_50_/mL at peak replication. The cells containing great roundleaf bat ACE2 had peak SC2 titers of 10^5.8^ TCID_50_/mL resulting in significantly higher viral titers then the rest of the bat-ACE2 cells. SC2 peaked around 10^5^-10^5.5^ TCID_50_/mL in most of the cells containing a bat ACE2, but the cells containing ACE2 from Egyptian rousette and Chinese rufous horseshoe bat grew significantly lower titers of the virus during the initial infection reaching only 10^4.7^ TCID_50_/mL and 10^4.4^ TCID_50_/mL, respectively (Figure 3A). However, after subsequent passage onto fresh cells, the Wuhan lineage isolate grew similarly in all cell lines, around 10^4^ TCID_50_/mL, except in the DF1 (+h/+h) cells, which grew SC2 roughly 2-logs higher (Figure 3B). As previously observed, the DF1 (-/-) cells expressing only chicken ACE2 did not become infected (data not shown) (30). Cytopathic effect (CPE) caused by the Wuhan lineage SC2 was observed in all cells except for DF1 (-/-), as expected (Figure 3C). SC2 S expression was detected using a mouse anti-SARS-CoV-2 S monoclonal antibody. The DF1 (+h/+h) cell line demonstrated the highest immunostaining, but S was detected in all cells expressing bat ACE2 indicating the presence of an infection (Figure 3C). These results indicate that the bat species tested here are permissible to the Wuhan lineage strain of SC2.

**Figure 3.**
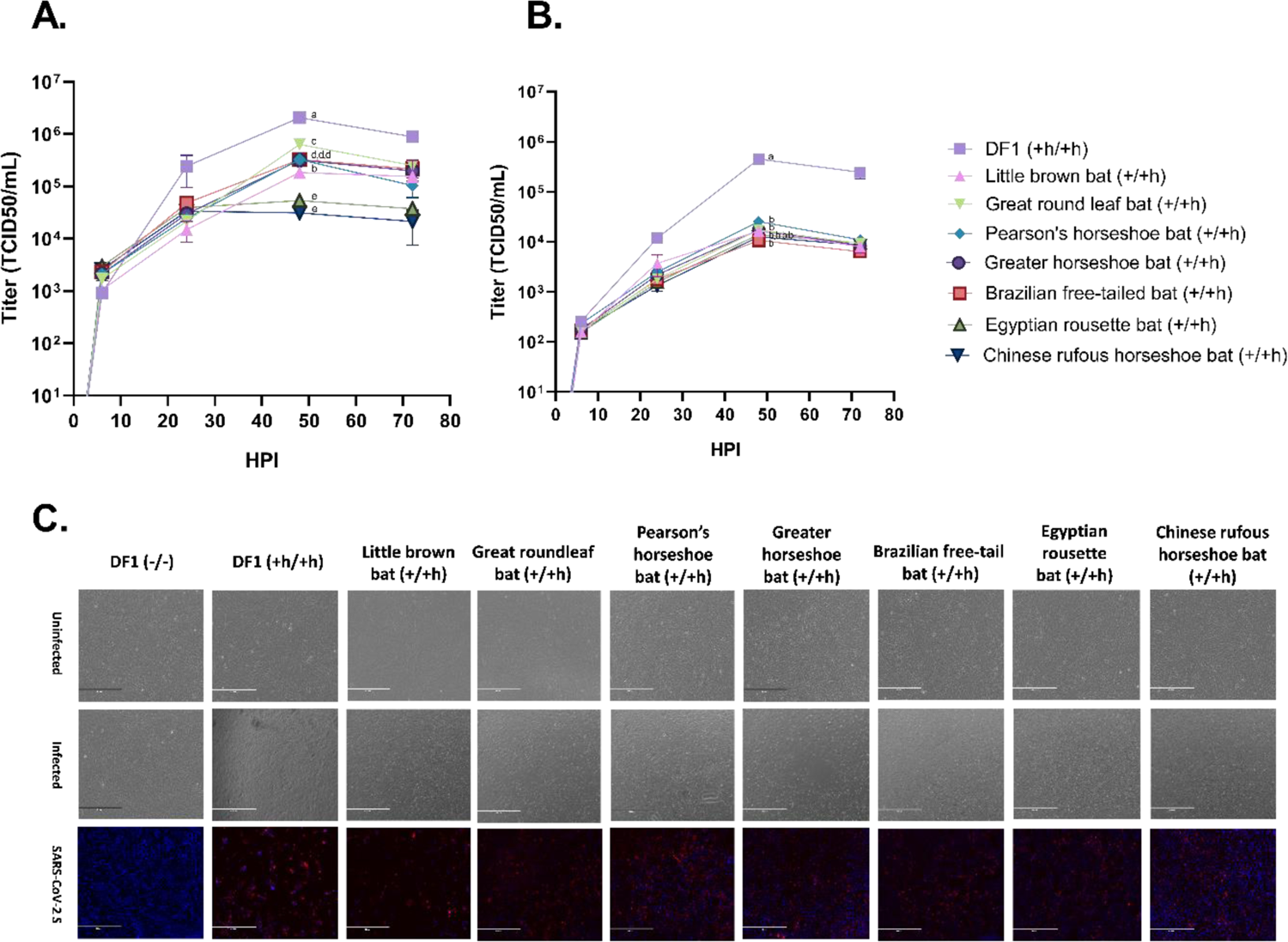
Growth of SARS-CoV-2 (Wuhan lineage) in DF1 cells expressing bat ACE2, human TMPRSS2. DF1 cells expressing bat ACE2 and human TMPRSS2 cells were infected with the Wuhan lineage strain of SC2 at an MOI of 1. At 2, 6, 24, 48, and 72 HPI supernatant samples were taken for RNA extraction, and viral titers were determined by RT-qPCR. The values shown are the mean with standard deviation of triplicate samples. Two-way analysis with Tukey’s multiple comparison test was performed on viral titers at 48 HPI to determine the statistical significance of viral titer between cell lines. Different lowercase letters indicate significant differences (p<0.05). **(A)** Pass 1 of the virus in cell culture. **(B)** Pass 2 of the virus in cell culture. After 72 HPI, supernatants from pass 1 were transferred onto fresh monolayers for 1 hour, washed with PBS, and replaced with fresh media. The time points from pass 1 were repeated. **(C)** DF1 cells expressing bat ACE2 and human TMPRSS2 were grown on glass chamber slides. Cells were infected at an MOI of 1. At 48 HPI, cells were imaged to examine CPE. Images of uninfected and infected cells were taken. Cells were also fixed and stained with a rabbit-anti-SARS-CoV-2-S antibody followed by a goat anti-rabbit Cy3-conjugated secondary antibody. Cells were counterstained with DAPI and visualized on an EVOS 500 microscope.

Growth of Delta SC2 was less variable in the cell lines expressing a bat species ACE2. All cell lines had peak Delta titers at 10^4^-10^4.3^ TCID_50_/mL apart from the DF1 (+h/+h) cell line, which had peak Delta titers similar to Wuhan-lineage SC2 (>10^6^ TCID_50_/mL) during the initial round of infection (Figure 4A). During pass 2 of Delta, the titers decreased to approximately 10^3^ TCID_50_/mL in the cell lines containing a bat ACE2 (Figure 4B). Although Delta titers were much lower in the cells containing a bat ACE2 than the DF1 (+h/+h) control, CPE and S immunostaining was observed in all cell lines (Figure 3C). The data indicate that although the Delta variant can infect all bat ACE2-expressing cell lines tested, replication may be less efficient than Wuhan lineage SC2 in this model.

**Figure 4.**
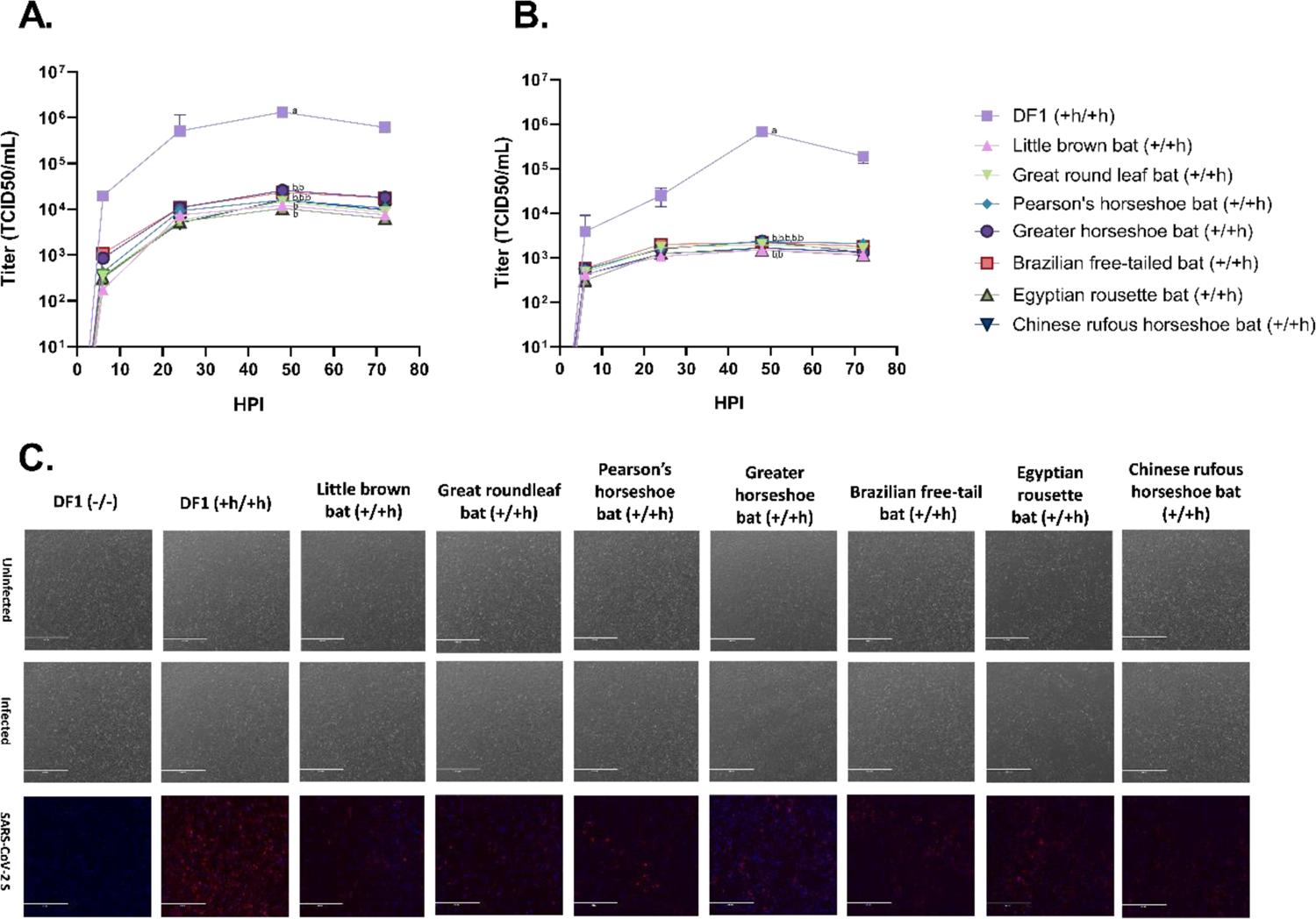
Growth of SARS-CoV-2 (Delta) in DF1 cells expressing bat ACE2, human TMPRSS2. DF1 cells expressing bat ACE2 and human TMPRSS2 cells were infected with the Delta strain of SC2 at an MOI of 1. At 2, 6, 24, 48, and 72 HPI supernatant samples were taken for RNA extraction, and viral titers were determined by RT-PCR. The values shown are the mean with standard deviation of triplicate samples. Two-way analysis with Tukey’s multiple comparison test was performed on viral titers at 48 HPI to determine the statistical significance of viral titer between cell lines. Different lowercase letters indicate significant differences (p<0.05). **(A)** Pass 1 of the virus in cell culture. **(B)** Pass 2 of the virus in cell culture. After 72 HPI, supernatants from pass 1 were transferred onto fresh monolayers for 1 hour, washed with PBS, and replaced with fresh media. The time points from pass 1 were repeated. **(C)** DF1 cells expressing bat ACE2 and human TMPRSS2 were grown on glass chamber slides. Cells were infected at an MOI of 1. At 48 HPI, cells were imaged to examine CPE. Images of uninfected and infected cells were taken. Cells were also fixed and stained with a rabbit-anti-SARS-CoV-2-S antibody followed by a goat anti-rabbit CY3-conjugated secondary antibody. Cells were counterstained with DAPI and visualized on an EVOS 500 microscope.

Analysis of Lambda SC2 variant demonstrated significantly lower replication in the bat ACE2 cell lines compared to the DF1 (+h/+h) control cells. However, variability of Lambda growth was greater in these lines. Lambda had peak titers at or above 10^4^ TCID_50_/mL in the greater horseshoe (10^4.4^ TCID_50_/mL), Brazilian free-tailed (10^4.5^ TCID_50_/mL), and Pearson’s horseshoe (10^4.3^ TCID_50_/mL) bat cells, whereas titers remained below 10^4^ TCID_50_/mL in the rest of the cells with bat ACE2 on pass 1 (Figure 5A). Lambda grew the DF1 (+h/+h) control cells comparably to Wuhan lineage and Delta SC2 at >10^6^ TCID_50_/mL. Consistent with the other two viruses tested, Lambda titers dropped on pass 2. Although the difference in Lambda growth was not significant between any of the bat ACE2 cell lines, the Brazilian free-tailed bat ACE2 cell line grew the virus to slightly higher titers, at 10^3.4^ TCID_50_/mL, than the other cell lines tested (Figure 5B). Cytopathic effect and S expression was observed in all cell lines except the negative-control cell line, DF1 (-/-) (Figure 5C). Taken together, the results demonstrate that Lambda can utilize the bat ACE2 for entry into DF1 cells, which were permissible to virus replication.

**Figure 5.**
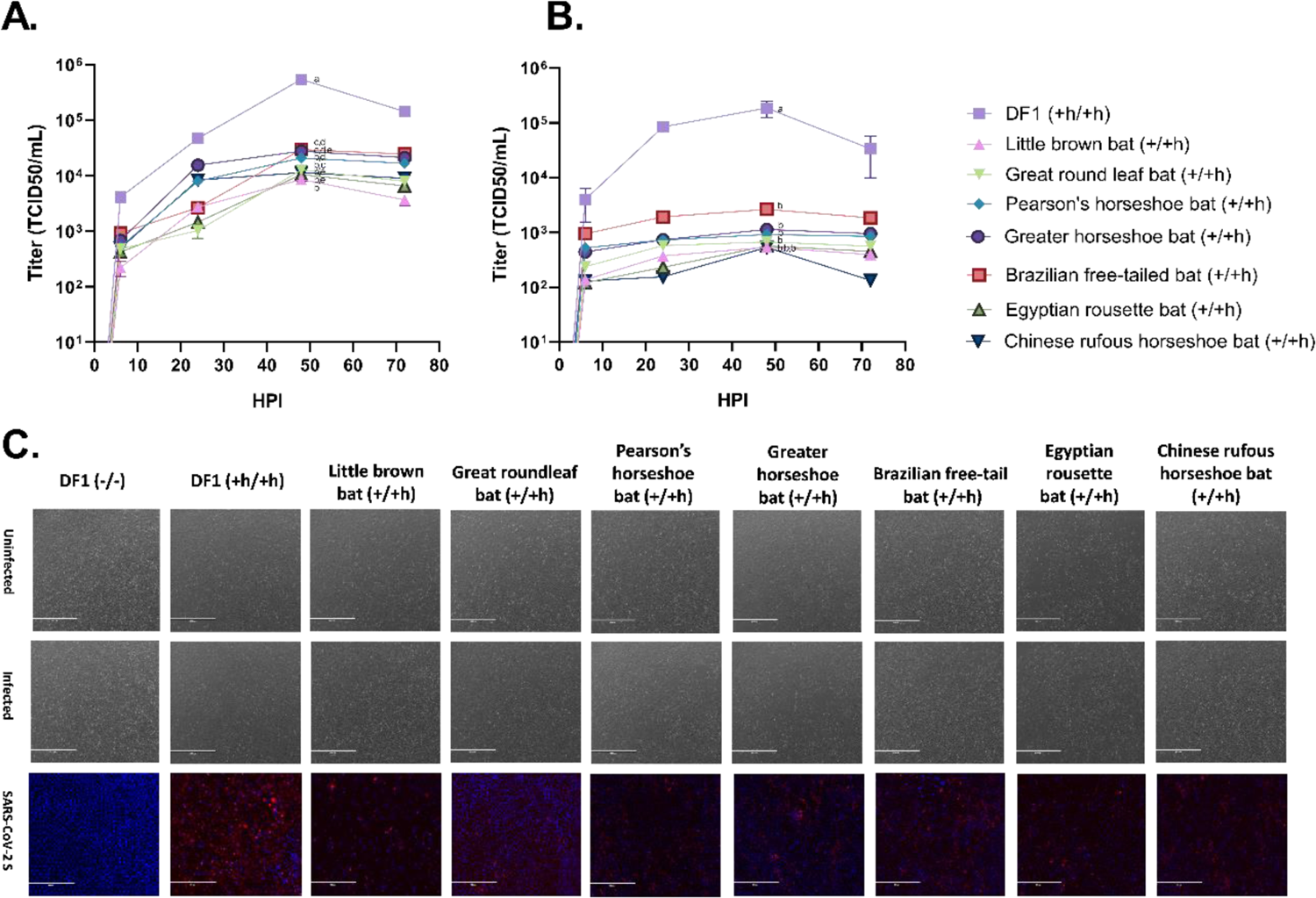
Growth of SARS-CoV-2 (Lambda) in DF1 cells expressing bat ACE2, human TMPRSS2. DF1 cells expressing bat ACE2 and human TMPRSS2 cells were infected with the Lambda strain of SC2 at an MOI of 1. At 2, 6, 24, 48, and 72 HPI supernatant samples were taken for RNA extraction, and viral titers were determined by RT-PCR. The values shown are the mean with standard deviation of triplicate samples. Two-way analysis with Tukey’s multiple comparison test was performed on viral titers at 48 HPI to determine the statistical significance of viral titer between cell lines. Different lowercase letters indicate significant differences (p<0.05). **(A)** Pass 1 of the virus in cell culture. **(B)** Pass 2 of the virus in cell culture. After 72 HPI, supernatants from pass 1 were transferred onto fresh monolayers for 1 hour, washed with PBS, and replaced with fresh media. The time points from pass 1 were repeated. **(C)** DF1 cells expressing bat ACE2 and human TMPRSS2 were grown on glass chamber slides. Cells were infected at an MOI of 1. At 48 HPI, cells were imaged to examine CPE. Images of uninfected and infected cells were taken. Cells were also fixed and stained with a rabbit-anti-SARS-CoV-2-S antibody followed by a goat anti-rabbit CY3-conjugated secondary antibody. Cells were counterstained with DAPI and visualized on a EVOS 500 microscope.

## DISCUSSION

Since 2019, much work has been done on SC2, but its origin remains unclear. Several established cell lines have been used to study virus growth (Vero, Caco-2, Calu-3, 293T), but most are naturally permissive to the virus and cannot be used for testing host susceptibility (41, 42). DF1 cells, however, have been shown to be non-permissive to SC2 and can act as a cellular backbone for testing various animal ACE2 and TMPRSS2 genes (39). We previously showed that DF1 cells expressing the ACE2 and TMPRSS2 genes from different animal species can be used as an *in vitro* predictive model for virus replication (34). Here we utilize the same method to examine host range and susceptibility in seven bat species.

In this study, we used human TMPRSS2 for all bat ACE2 transgenic cells because of the inconsistent availability and reliability of bat TMPRSS2 sequences. In our previous study, we observed that the little brown bat and great round leaf bat cell lines could bind the virus, but only transient viral replication was observed within them. We suspected this was due to wrongly annotated TMPRSS2 genes in GenBank, which resulted in a partial sequence for both species (34). To circumvent this problem and the lack of TMPRSS2 sequences available for some bat species, human TMPRSS2 was used. We postulated that human TMPRSS2 could be used as a substitute because the amino acids in the active site are conserved between bats and humans. Further research is underway to determine the correct TMPRSS2 sequences for several bat species.

The DF1 (+h/+h) cell line was previously developed using a lentivirus vector to insert both ACE2 and TMPRSS2, whereas the DF1 cells containing the bat ACE2 gene were developed using a transposon vector system. The lentivirus appears to be a more efficient delivery system and resulted in higher expression of ACE2 and TMPRSS2 in the DF1 (+h/+h) cells. Additionally, TMPRSS2 was inserted first in the transgenic cell lines resulting in higher expression compared to ACE2 (Figure 2 and Supplemental figure 3). We cannot exclude the possibility that the differences in ACE2 expression led to the differences in viral replication efficiency. However, we demonstrated that the bat-origin ACE2 tested here are viable receptors for three variants of SC2 and can infer the likely ability of these bat species to become infected and potentially act as a vector for virus transmission to other species.

In this study, we expanded previous work looking at host susceptibility by examining seven different bat species from different parts of the world. The bat species have different ranges, roosting sites, and foraging habits, which affect the amount of contact they have with humans. For example, Egyptian rousettes, Brazilian free-tailed bats, and little brown bats have increased human contact compared to the horseshoe bat species, which tend to live in more rural areas (41). Of particular interest for this study were the *Rhinolophus* (horseshoe) species (Pearson’s horseshoe bat, greater horseshoe bat, and Chinese rufous horseshoe bat), which are known to carry other bat coronaviruses, and are thought to be a possible host for SC2 (25). Our study shows that all three species of horseshoe bat ACE2 were able to support entry and viral replication of all three variants of SC2 (Figures 3-5). To the best of our knowledge, no *in vivo* testing has occurred in a *Rhinolophus* species, but a large study using a SC2 pseudodovirus examined viral entry into cells of various bat species. The study used 293T cells transduced with different bat ACE2 orthologues, but did not transduce TMPRSS2, which is required for increased infectivity and viral replication (34, 41). They found that infection efficiency was <5% for Pearson’s horseshoe bat, greater horseshoe bat, and Chinese rufous horseshoe bat ACE2, and surmised viral replication could not be supported in these species, which greatly differs from our findings. Additionally, great round leaf bat ACE2, which is highly similar to that of the *Rhinolophus* species, did not support SC2 entry in their assay. We found that SC2 growth was supported using our great round leaf bat ACE2 cell line. Egyptian rousettes, Brazilian free-tailed bats, and little brown bats were found to support SC2 entry in their study, which our findings also support (Figures 3-5) (41).

Egyptian rousettes were experimentally infected with Wuhan lineage SC2 and found to have prominent viral titers in their respiratory tract and developed neutralizing antibodies (28). Our Egyptian rousette ACE2 cells replicated the viruses in a similar manner, but Delta and Lambda had decreased titers at peak replication (Figures 3-5). We observed an overall decrease in Delta and Lambda viral titers in the cells with a bat ACE2, and we postulate this is due to those variants being more human adapted than the Wuhan lineage. More recently, two studies looking at the susceptibility of the Brazilian free-tailed bat showed that the species can become infected (without showing symptoms) and develop antibodies to SC2. However, both studies found no evidence of viral transmission to uninfected bats (26, 27). Our results correlate with these findings as the Brazilian free-tailed bat ACE2 cell line was able to replicate all three variants of SC2 (Figures 3-5). Interestingly, big brown bat (*Eptesicus fuscus*) was found to be resistant to an infectious challenge with SC2 (29). Big brown bat susceptibility or that of a highly similar bat species has yet to be tested in our model.

Predictive *in silico* analyses of animal ACE2 sequences provide limited knowledge about species susceptibility, and *in vivo* SC2 challenges with wild animals present numerous logistical and ethical issues. The development of *in vitro* assays has been essential in determining SC2 susceptibility and host range. Here we expand on a previously reported rapid and economical method to screen susceptibility of ACE2 from seven bat species to three variants of SC2 (34).

## Acknowledgements

We thank Scott Lee, Suzanne DeBlois, and Tim Olivier for excellent technical assistance. This research was supported by funding from U.S. Fish and Wildlife Service agreement # 4500148748; USDA, APHIS, VS IAA # 22-9206-0593-IA; and USDA, ARS, CRIS project #6040-32000-081-00D. The USA/WA1/2020/Wuhan lineage (BEI NR-52286; Wuhan lineage strain), USA/PHC658/2021/B.1.617.2 (BEI NR-55611; Delta strain), and Peru/un-CDC-2-4069945/2021/Lineage C.37 (BEI NR-55654; Lambda strain) of SARS-CoV-2 was obtained from BEI Research Resources Repository, National Institute of Allergy and Infectious Diseases, National Institutes of Health. Vero African Green Monkey Kidney Cells (ATCC® CCL-81™), FR-243, were obtained through the International Reagent Resource, Influenza Division, WHO Collaborating Center for Surveillance, Epidemiology and Control of Influenza, Centers for Disease Control and Prevention, Atlanta, Georgia, USA. Any use of trade, firm, or product names is for descriptive purposes only and does not imply endorsement by the U.S. Government.

## Data availability

The data are available from the senior author upon request.

