## Supplemental data for "SARS-CoV-2 utilization of ACE2 from different bat species allows for virus entry and replication *in vitro*"

**Supplemental Figure 1**. Sequence similarity of angiotensin-converting enzyme 2 (ACE2) proteins. Sequences were obtained from Genbank and aligned using Geneious Prime[1]. A global alignment with free end gaps was performed on the ACE2 sequences. Table represents percent similarity (Blosum45 with threshold 0) based on protein sequence.

**Supplemental Figure S2. Alignment of human and bat ACE2 proteins.** Amino acid comparison of ACE2 examined. ACE2 domains are denoted in maroon, and the signal peptide is in pink. The legend for percent similarity is provided. Alignment was made using Geneious Prime. A global alignment with free end gaps was performed on ACE2 protein sequences.

**Supplemental Figure S3: Expression of TMPRSS2 in DF1 (avian fibroblast) cells.** DF1 cells lines containing bat ACE2 and human transmembrane serine protease 2 (TMPRSS2) were examined for expression of ACE2. **(A)** TMPRSS2 mRNA levels were measured in the cell lines using RT-qPCR. Values were normalized to chicken 28S. DF1 (-/-) cells were used as a negative control to calculate the ΔΔCT values. **(B)** human TMPRSS2 protein expression was measured by western blot. Cell lysates were resolved by SDS-PAGE and then transferred to a polyvinylidene difluoride (PVDF) membrane. The blot was probed with a rabbit anti-human TMPRSS2 and a rabbit anti-human actin antibody, followed by a goat anti-rabbit Dylight™ 594 secondary antibody for detection.

[1] USGS Disclaimer: Any use of trade, firm, or product names is for the descriptive purposes only and does not imply endorsement by the U.S. Government.

### Figure S1

| | Human | Little brown bat | Great roundleaf bat | Pearson's horseshoe bat | Greater horseshoe bat | Brazil free-tailed bat | Egyptian rousette bat | Chinese rufous horseshoe bat | Golden hamster | Goat | Cat | Pig | Horse | Chicken |
|---|---|---|---|---|---|---|---|---|---|---|---|---|---|---|
| Human | | 92.81 | 93.92 | 94.91 | 94.41 | 93.79 | 93.54 | 94.16 | 96.02 | 94.41 | 95.65 | 94.04 | 96.15 | 88.02 |
| Little brown bat | 92.81 | | 91.97 | 92.33 | 92.2 | 93.17 | 91.35 | 92.2 | 92.94 | 92.2 | 93.06 | 92.57 | 93.79 | 86.06 |
| Great roundleaf bat | 93.92 | 91.97 | | 94.67 | 95.04 | 93.3 | 92.68 | 94.29 | 94.04 | 92.06 | 94.29 | 92.56 | 94.42 | 87.55 |
| Pearson's horseshoe bat | 94.91 | 92.33 | 94.67 | | 97.27 | 93.91 | 93.91 | 98.39 | 94.41 | 93.42 | 95.78 | 93.04 | 94.78 | 87.41 |
| Greater horseshoe bat | 94.41 | 92.2 | 95.04 | 97.27 | | 93.29 | 94.29 | 97.14 | 94.53 | 93.79 | 95.4 | 93.54 | 95.4 | 87.16 |
| Brazil free-tailed bat | 93.79 | 93.17 | 93.3 | 93.91 | 93.29 | | 92.92 | 93.91 | 92.92 | 92.67 | 95.03 | 93.29 | 94.04 | 86.91 |
| Egyptian rousette bat | 93.54 | 91.35 | 92.68 | 93.91 | 94.29 | 92.92 | | 93.79 | 93.42 | 93.91 | 93.91 | 93.17 | 94.66 | 85.93 |
| Chinese rufous horseshoe bat | 94.16 | 92.2 | 94.29 | 98.39 | 97.14 | 93.91 | 93.79 | | 93.91 | 93.17 | 95.16 | 93.17 | 94.66 | 87.41 |
| Golden hamster | 96.02 | 92.94 | 94.04 | 94.41 | 94.53 | 92.92 | 93.42 | 93.91 | | 94.66 | 95.4 | 94.29 | 96.02 | 87.16 |
| Goat | 94.41 | 92.2 | 92.06 | 93.42 | 93.79 | 92.67 | 93.91 | 93.17 | 94.66 | | 94.91 | 95.78 | 95.4 | 86.79 |
| Cat | 95.65 | 93.06 | 94.29 | 95.78 | 95.4 | 95.03 | 93.91 | 95.16 | 95.4 | 94.91 | | 95.28 | 97.14 | 88.64 |
| Pig | 94.04 | 92.57 | 92.56 | 93.04 | 93.54 | 93.29 | 93.17 | 93.17 | 94.29 | 95.78 | 95.28 | | 95.78 | 87.28 |
| Horse | 96.15 | 93.79 | 94.42 | 94.78 | 95.4 | 94.04 | 94.66 | 94.66 | 96.02 | 95.4 | 97.14 | 95.78 | | 87.65 |
| Chicken | 88.02 | 86.06 | 87.55 | 87.41 | 87.16 | 86.91 | 85.93 | 87.41 | 87.16 | 86.79 | 88.64 | 87.28 | 87.65 | |

**Figure S2**

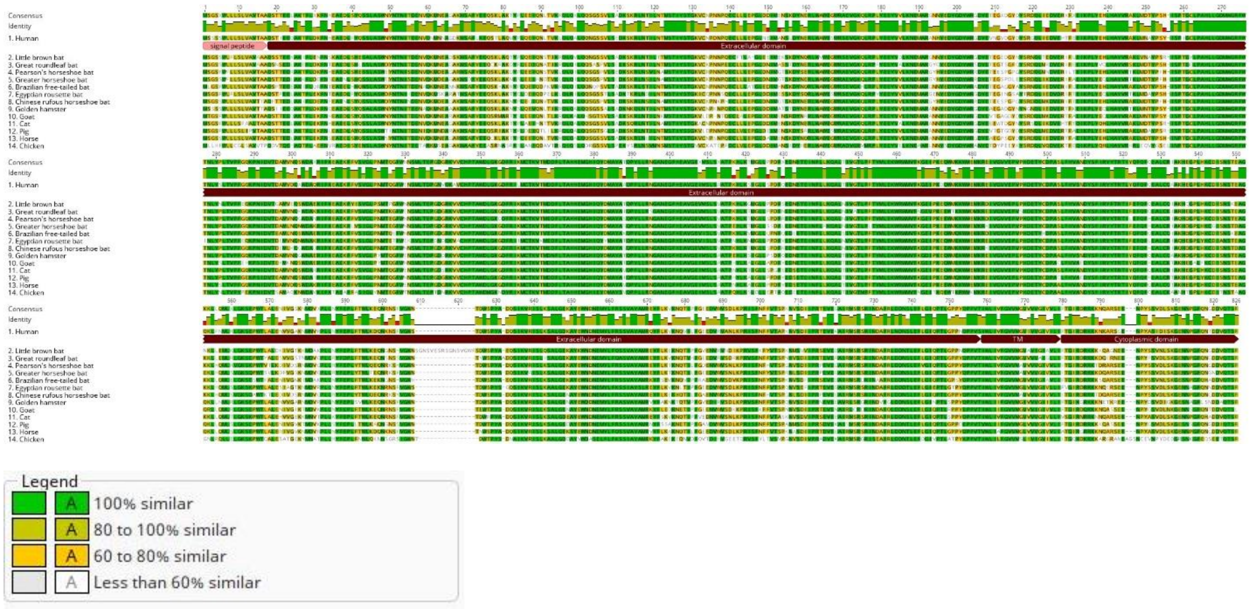
Consensus
Identity
1. Human
2. Little brown bat
3. Great roundleaf bat
4. Pearson's horseshoe bat
5. Greater horseshoe bat
6. Brazilian free-tailed bat
7. Egyptian rousette bat
8. Chinese rufous horseshoe bat
9. Golden hamster
10. Goat
11. Cat
12. Pig
13. Horse
14. Chicken
signal peptide
Extracellular domain
TM
Cytoplasmic domain
Legend
100% similar
80 to 100% similar
60 to 80% similar
Less than 60% similar

**Figure S3**

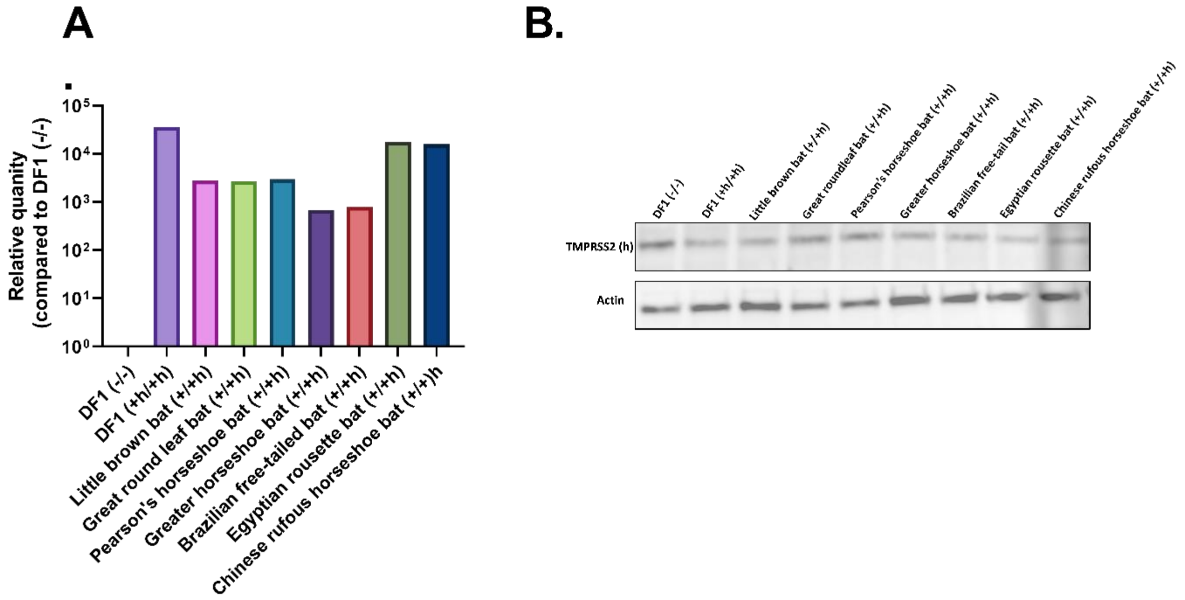
A
.
Relative quanity
(compared to DF1 (-/-)
$10^5$
$10^4$
$10^3$
$10^2$
$10^1$
$10^0$
DF1 (-/-)
DF1 (+h/+h)
Little brown bat (+/+h)
Great round leaf bat (+/+h)
Pearson's horseshoe bat (+/+h)
Greater horseshoe bat (+/+h)
Brazilian free-tailed bat (+/+h)
Egyptian rousette bat (+/+h)
Chinese rufous horseshoe bat (+/+)h
B.
DF1 (-/-)
DF1 (+h/+h)
Little brown bat (+/+h)
Great roundleaf bat (+/+h)
Pearson's horseshoe bat (+/+h)
Greater horseshoe bat (+/+h)
Brazilian free-tail bat (+/+h)
Egyptian rousette bat (+/+h)
Chinese rufous horseshoe bat (+/+h)
TMPRSS2 (h)
Actin
